# Accelerating cryptic pocket discovery using AlphaFold

**DOI:** 10.1101/2022.11.23.517577

**Authors:** Artur Meller, Soumendranath Bhakat, Shahlo Solieva, Gregory R. Bowman

## Abstract

Cryptic pockets, or pockets absent in ligand-free, experimentally determined structures, hold great potential as drug targets. However, cryptic pocket opening is often beyond the reach of conventional biomolecular simulations because certain cryptic pocket openings involve slow motions. Here, we investigate whether AlphaFold can be used to accelerate cryptic pocket discovery either by generating structures with open pockets directly or generating structures with partially open pockets that can be used as starting points for simulations. We use AlphaFold to generate ensembles for 10 known cryptic pocket examples, including 5 that were deposited after AlphaFold’s training data was extracted from the PDB. We find that in 6 out of 10 cases AlphaFold samples the open state. For plasmepsin II, an aspartic protease from the causative agent of malaria, AlphaFold only captures partial pocket opening. As a result, we ran simulations from an ensemble of AlphaFold-generated structures and show that this strategy samples cryptic pocket opening, even though an equivalent amount of simulations launched from a ligand-free experimental structure fails to do so. Markov state models (MSMs) constructed from the AlphaFold-seeded simulations quickly yield a free energy landscape of cryptic pocket opening that is in good agreement with the same landscape generated with well-tempered metadynamics. Taken together, our results demonstrate that AlphaFold has a useful role to play in cryptic pocket discovery but that many cryptic pockets may remain difficult to sample using AlphaFold alone.

## Introduction

Cryptic pockets, or pockets absent in ligand-free experimental structures, are a promising means to expand the scope of drug discovery. By one estimate, almost half of all structured domains lack obvious pockets in their experimental structures^1^. These proteins have often been considered ‘undruggable’. However, as proteins fluctuate in solution, they may adopt excited structural states that contain cryptic pockets. Thus, cryptic pockets may provide a means to target these ‘undruggable’ proteins^2^. Furthermore, many cryptic pockets are distant from active sites, suggesting that targeting them may lead to the discovery of allosteric activators^3^ or more specific modulators given the high sequence conservation of many active sites^4^.

The discovery of cryptic pockets using experimental and computational methods remains difficult in many cases. Most cryptic pockets are discovered serendipitously when experimental structures of a ligand bound to a protein reveal a novel binding site that is closed in ligand-free structures of the same protein^5^ While this process has revealed cryptic pockets, it requires knowledge of a ligand *a priori*. Molecular dynamics simulations can reveal excited states with cryptic pockets that can then be used for structure-based drug design^2,6^. However, in certain cases, cryptic pockets may not be discovered by simulations because cryptic pocket opening motions may be slow (e.g., Niemann-Pick C2 Protein in Meller et al.^1^). Two classes of slow motions include sidechain ring flipping^7^ events and secondary structure rearrangements^8^ which can both occur on microsecond and slower timescales.

Here we explore the possibility of using AlphaFold^9^ to accelerate cryptic pocket discovery. Previous work has shown that stochastic sampling of AlphaFold’s input multiple sequence alignment can generate diverse conformations of membrane and globular proteins^10,11^. We hypothesized that a similar strategy can be applied to discover cryptic pockets. Even if AlphaFold can only capture partial opening, we reasoned that starting molecular dynamics simulations from these structures may capture full opening far more quickly than starting simulations from completely closed structures.

We test our strategy of launching simulations from AlphaFold-generated starting structures with plasmepsin II (PM II), a well-studied protease from the causative agent of malaria^12–14^. PM II is one of many aspartic proteases that play an important role in the lifecycle of *Plasmodium falciparum*. It is found in digestive vacuoles where it is used by the parasite to digest hemoglobin. Though functional redundancy in digestive vacuoles may limit the utility of narrow PM II inhibitors, PM II may play a role in antimalarial drug resistance^15^ and provide insight into developing inhibitors of other aspartic proteases that are essential in the Plasmodium lifecycle. Notably, PM II contains a cryptic pocket adjacent to its active site, which was revealed in several experimental structures capturing PM II bound to different classes of inhibitors. Given that previous simulation studies of PM II have failed to sample cryptic pocket opening^13^, here we explore if increasing aggregate simulation time is sufficient to open this pocket or if AlphaFold can accelerate cryptic pocket discovery.

## Methods

### Ensemble generation using AlphaFold

To generate ensembles of structures from a sequence rather than a single structure, we use two modifications to the original AlphaFold implementation. Firstly, we stochastically subsample the multiple sequence alignment (MSA) to a maximum of 32 cluster centers and 64 extra sequences. Each time we generate a structure prediction a different random seed is used for sequence clustering, so that the input MSA passed to AlphaFold is slightly modified. Secondly, we also enable dropout during the forward-pass through the model.

We generated ensembles for each of the proteins studied using ColabFold^16^, a fast and userfriendly implementation of the AlphaFold algorithm. Specifically, we used the Google Collaboratory notebook. We generated initial MSAs using the *jackhammer* method with pre-filtering that enforced a minimum 50% coverage and 20% sequence identity with the query. We then limited the depth of the input MSA by setting the *max_msa_clusters* variable to 32 and *max_extra_msa* to 64. We generated ensemble of 32 or 160 structures by setting *num_models* to 1 or 5 respectively and *num_samples* to 32. We enabled *droupout* by setting *is_training* to True. We also enabled *use_ptm*, set *num_ensembles* to 1, set *tol* to 0, and set *max_recycles* to 3.

The link to the Google Collaboratory notebook is here (https://colab.research.google.com/github/sokrypton/ColabFold/blob/main/beta/AlphaFold2_advanced.ipynb).

### Molecular dynamics simulations

We prepared molecular dynamics simulations using the tleap module integrated with Amber 2020^17^ with the workflow described here. Proteins were parametrized using the AMBER FF14SB^18^ force field and solvated in a truncated octahedron box with TIP3P^19^ waters. Each system was neutralized by 17 Na^+^ ions. For each system, the box was extended 1.0 nm from protein atoms in all directions. Minimization was performed in two steps: (a) initial minimization where the protein was constrained with a restrained potential of 100 kcal/mol^-1^Å^2^ to minimize only the water and ions (200 steps of steepest descent followed by 200 steps of conjugate gradients) followed by (b) 500 steps of unrestrained minimization of the whole system.

We equilibrated protein systems and performed production runs using Gromacs 2021^20^. Following minimization in Amber, we converted Amber topologies to Gromacs format using Acpype^21^. Initially, we heated each system (from 0 K to 300 K) using the NVT ensemble for 500 ps with harmonic restraints of 500 kJ mol^-1^nm^-2^ applied to backbone heavy atoms. Next, each system was equilibrated at 300 K in an NPT ensemble for 200 ps without any restraints using the Parrinello-Rahman barostat^22^ to maintain the pressure at 1 bar and the v-rescale thermostat for temperature control. Production runs were carried out in the NPT ensemble at 300 K and 1 bar using the leap-frog integrator and Parrinello-Rahman thermostat with a 2 fs timestep. Non-bonded interactions were cut off at 1.0 nm, and long-range electrostatic potentials were treated using the Particle Mesh Ewald (PME) method^23^ with a grid spacing of 0.16 nm. The LINCS algorithm^24^ was used to constrain H-bonds during MD simulations.

We performed 640 independent MD simulations in total to generate apo-seeded and AF-seeded ensembles each with 32 microseconds of sampling. We used 32 different AlphaFold-generated starting structures for plasmepsin II with 10 independent (i.e., starting from different initial velocities) simulations launched for each structure. Each simulation was 100 ns in length. For the apo-seeded ensemble, we ran 320 independent simulations 100 ns in length starting from a single starting structure from the PDB (1LF4^25^).

### Markov State Modeling

To construct MSMs^26–28^, we first defined a subset of features that were relevant to PM II cryptic pocket opening. We focused on the set of residues that were within 0.5 nm of the cryptic extension of the A1T ligand in the holo crystal structure (PDB: 2IGX^29^). Specifically, we located all residues that were within 0.5 nm of the following A1T atoms: C48, C46, C43, C40, C38, C36, C33, C34, C30, N29, and C26. We then used backbone (phi, psi) and sidechain dihedrals for those residues to define an initial feature set relevant for cryptic pocket opening. We removed any χ-2 angles that included symmetrically equivalent atoms (e.g., χ-2 for tyrosine residues).

To perform clustering in a kinetically relevant space, we applied time-structure-independent component analysis^30^ (tICA) to these features. Specifically, we used a tICA lag time of 10 ns and retained the top *n* tICs that accounted for 90% of kinetic variance using commute mapping.

To determine the appropriate number of microstates for clustering, we used a cross-validation scheme where trajectories were partitioned into training and test sets. Clustering into k microstates was performed using only the training set, and the test set trajectories were assigned to these k microstates based on their Euclidean proximity in tICA space to each microstate’s centroid. Using the test set only, an MSM was fit using maximum likelihood estimation (MLE), and the quality of the MSM was assessed with the rank-10 VAMP-2 score of the transition matrix. We found that 25 microstates had the highest VAMP-2 score on average across 10 trials on the test set for the AF-seeded ensemble (Fig. S17). For consistency, we used the same number of microstates for the apo-seeded MSM.

Finally, MSMs of the PM II cryptic pocket were fit for the apo-seeded and AF-seeded ensembles separately using MLE. Lag times were chosen by the logarithmic convergence of the implied timescales test (Fig. S18, S19). Lag times of 12.5 ns were used for both the apo-seeded and AF-seeded MSMs.

MSM construction was performed using the PyEMMA^31^ software package.

### Metadynamics

We performed well-tempered metadynamics^32,33^ (WTMeta) simulations to sample the conformational landscape associated with Trp41 ring flipping, one of the motions necessary for plasmepsin II cryptic pocket opening. For each residue, we performed two-dimensional WTMeta at 300 K using χ-1 and χ-2 angles as collective variables. Gaussians were deposited every 500 time steps with a width and height of 0.05 radians and 1.2 kJ/mol respectively and a bias factor of 20. Unbiased free energy surfaces along different collective variables were extracted from WTMeta using the reweighting protocol described by Tiwary and Parrinello^34^.

We also used WTMeta simulations to study unbinding of small molecules from two *holo* conformations (PBD: 2BJU^35^, 4AY8^36^). Small molecules were parameterized using the General Amber Force Field^37^ (GAFF) and the protein was parameterized using the Amber14SB force field. The complexes were neutralized using sodium ions and immersed into a truncated octahedral box such that the distance from protein to the edge of the box was at least 1 nm. Equilibration and production runs were performed using the protocol described in Bhakat & Soderhjelm^13^. To estimate the apparent free energy profile of ligand unbinding, we performed multiple independent WTMeta simulations using the distance between the center of mass of the active side residues and the ligand as collective variables. All unbinding WTMeta simulations were performed at 300 K with a bias factor of 10 using Gaussian width and height of 0.011 nm and 1.2 kJ/mol respectively.

## Results

### AlphaFold predicts some but not all known cryptic pocket openings

We reasoned that AlphaFold (AF) could produce conformations with open cryptic pockets through stochastic sampling of its input multiple sequence alignment. Previous studies have shown that AlphaFold samples diverse conformations of transporters and receptors when its input MSA is stochastically subsampled to only include 16 sequences^10,11^. Additionally, AF ensembles of a set of proteins where ligand binding is associated with conformational rearrangements (though not necessarily at the ligand binding site) often included *holo-like* conformations^38^. However, it was not known if AlphaFold samples open structures for proteins known to form cryptic pockets when bound to drug-like molecules (e.g., not ions).

We generated AlphaFold ensembles for 10 known cryptic pocket examples, including a subset that was deposited to the PDB after AlphaFold was trained. These examples include several different types of conformational rearrangements: loop motions, secondary structure motions, and interdomain motions. To ensure that the network was not ‘memorizing’ particular conformations in its training dataset, we also focused on 5 cryptic pocket examples that were deposited to the PDB after April 2018, the date when the AlphaFold training set was pulled. We used ColabFold’s implementation of AlphaFold to generate 160 conformers for each input sequence because it offered a massive speed up and supported stochastic clustering of the input MSA (see Methods). We also used dropout in the forward-pass through the network to amplify structural diversity.

We find that AlphaFold samples many but not all cryptic pocket openings (Fig. 2). Among proteins that were in the training dataset, AlphaFold recapitulates known cryptic pockets in 3 out of 5 examples. In those cases, AF predicts a structure with less than 1.2 Å root mean square deviation (RMSD) to the *holo* structure in the cryptic site (i.e., using all heavy atoms within 5 Å of where the cryptic ligand binds for the RMSD calculation). Interestingly, AlphaFold generates open states of the Niemann-Pick C2 Protein that were not discovered in 2 microseconds of adaptive sampling simulations (Fig. 2B)^1^. However, AlphaFold’s ensemble of TEM β-lactamase structures does not include any open states where the Horn^39^ or omega^6^ pockets are open (Fig. S1). Among proteins that were not in the training dataset, AlphaFold recapitulates 3 of the 5 cryptic pockets (i.e., using pocket RMSD of 1.2 Å as the cutoff again). There appears to be a correlation between the size of the rearrangement (i.e., RMSD between *apo* and *holo* structures) and the ability of AF to sample cryptic pockets (Fig. S2-S12). For example, cryptic pocket opening in fascin requires a large interdomain motion (0.47 pocket RMSD between *apo* and *holo*) and is not captured in the AF ensemble.

**Figure 1.**
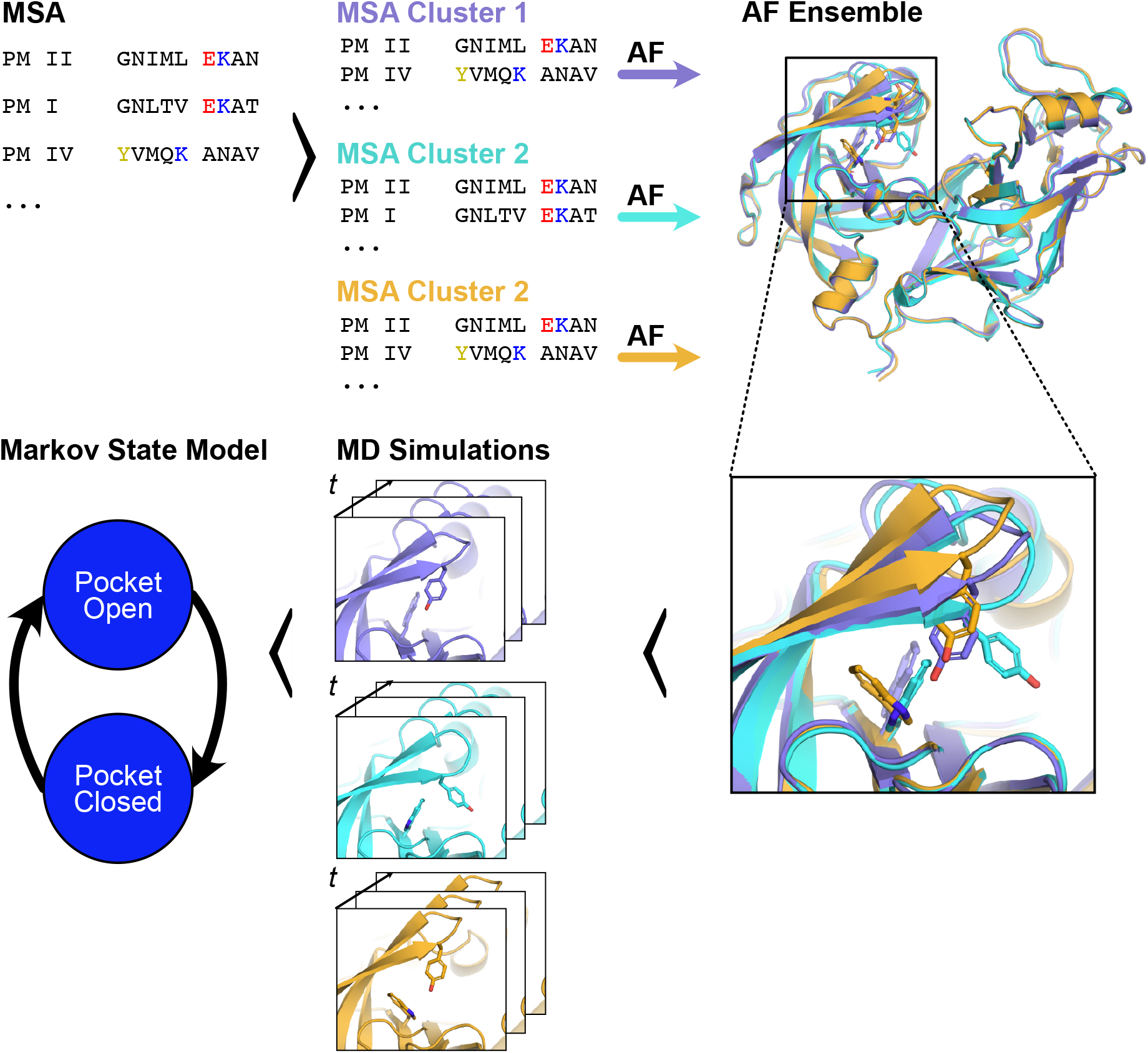
To efficiently sample cryptic pocket opening, we propose launching molecular dynamics simulations from diverse AlphaFold-generated starting conformations. Starting with an MSA of a query sequence (*top left*), the MSA can be stochastically clustered to create input MSAs of lower depth that are then fed to AlphaFold. Through this procedure, we can generate an ensemble of structures of the same protein (*top right* shows snapshots of Plasmodium falciparum’s plasmepsin II). These structures may adopt different conformations at known cryptic pockets (*bottom right* inset highlights different conformations of the plasmepsin II cryptic pocket). To generate free energy landscapes of cryptic pocket opening, we can launch molecular dynamics simulations from these different conformations and then stitch these simulations together with a Markov State Model.

**Figure 2.**
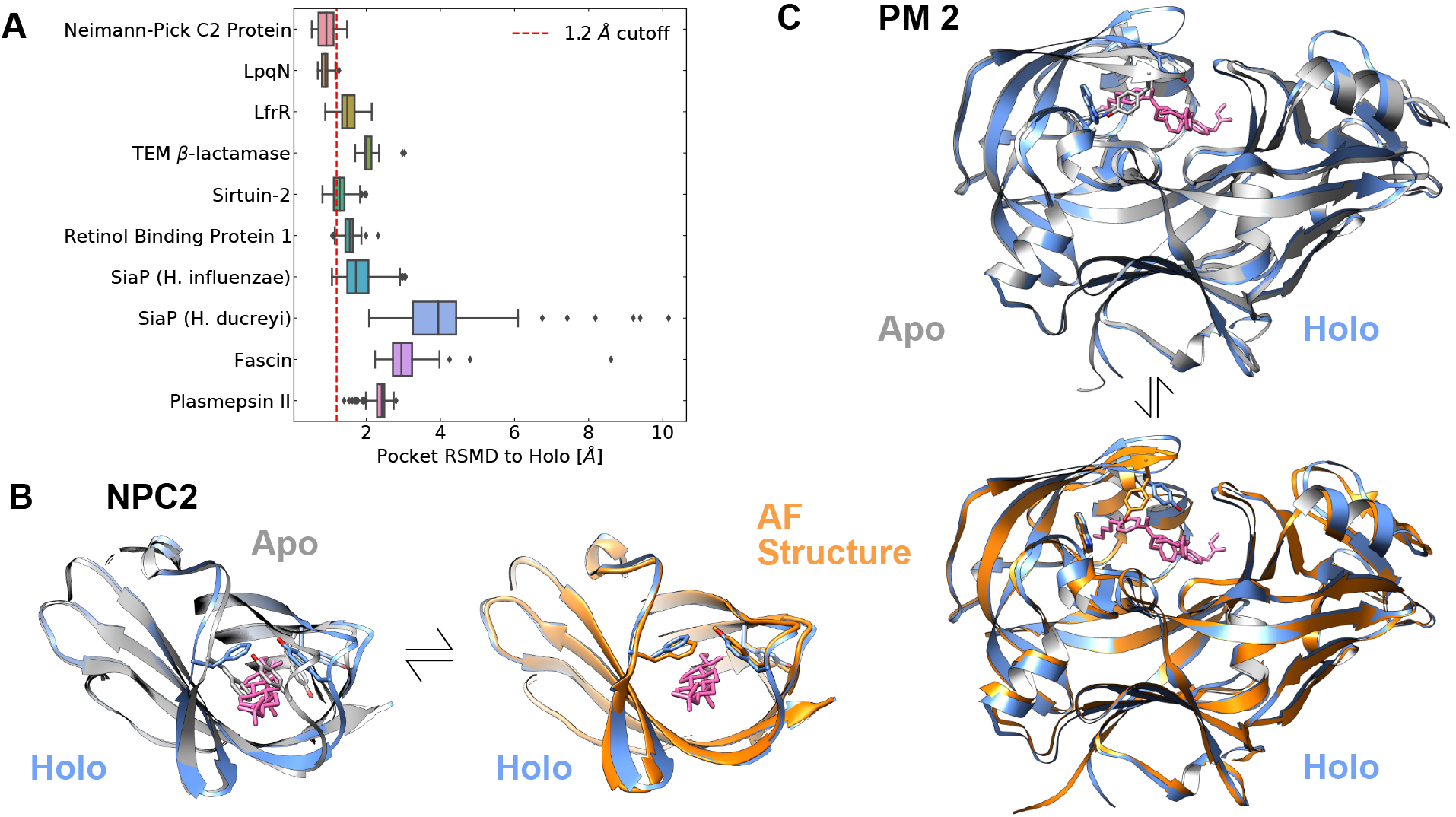
Stochastic clustering of its input multiple sequence alignment allows AlphaFold to generate structures with open or partially open cryptic pockets across multiple systems. **A)** In 6 out of 10 examples, AlphaFold samples the open state of a known cryptic pocket. The box-and-whisker plots show cryptic pocket root mean square deviation (RMSD) to a *holo* crystal structure (defined by heavy atoms within 5 Å of the ligand that binds at the cryptic pocket). For the top 5 examples, the *holo* structure was part of the training dataset for AlphaFold but the bottom 5 examples had their *holo* crystal structures deposited after AlphaFold was trained. The red line indicates 1.2 Å RMSD, a proposed cutoff for sampling the open state. **B)** Structural overlay of an AlphaFold-generated structure with the *holo* structure of Neimann-Pick C2 Protein (NPC2) shows that AlphaFold samples the open state. The ligand which binds in the cryptic pocket is shown in magenta; the *apo* structure is shown in gray; the *holo* structure is shown in blue; and the AF structure is shown in orange. Residues that change rotamer state between *apo* and *holo* experimental structures are shown in sticks. **C)** Structural overlay of an AlphaFold-generated structure of plasmepsin II with a *holo* structure containing a cryptic pocket shows that AlphaFold partially samples cryptic pocket opening. Select residues that change rotamer state between *apo* and *holo* experimental structures show that AlphaFold samples *holo*-like tryptophan orientations in the plasmepsin II cryptic pocket. As in B, the ligand which binds in the cryptic pocket is shown in magenta; the *apo* structure is shown in gray; the *holo* structure is shown in blue; and the AF structure is shown in orange.

Interestingly, for plasmepsin II (PM II), AlphaFold only samples partial cryptic pocket opening, capturing a ring flip that is necessary but not sufficient for pocket opening. In an AF-generated ensemble of 32 structures, there are several different Trp41 orientations (Fig. S13A). Notably, ligand-free PM II structures have only ever been observed in a single Trp41 orientation that blocks access to the cryptic site (Fig. S14). In contrast, the AF ensemble contains a Trp41 orientation that has only been experimentally observed in *holo* PM II structures with an open cryptic pocket (Fig. 2C, PDB: 2BJU^35^, 2IGX, 2IGY^29^). Similarly, AF-generated structures sample the Tyr77 conformation seen in *holo* PM II structures (Fig. S15). Despite this progress towards observing pocket opening, there are still significant differences in the position of the flap domain in the AF ensemble as compared to the *holo* crystal structures. In the AF ensemble, the flap domain has not moved away from the active site, sterically blocking known cryptic pocket binders. We wondered if simulations launched from the AF ensemble would sample cryptic pocket opening.

### PM II’s cryptic pocket opening is not captured with conventional MD simulations

We wanted to set a baseline to determine if AlphaFold accelerates cryptic pocket opening. Given recent success in using molecular dynamics to reveal cryptic pockets, we wondered if simulations launched from a ligand-free PM II structure would sample cryptic pocket opening. Though a previous study did not report cryptic pocket opening, it was limited to ~2 microseconds of sampling^13^. We hypothesized that increasing the aggregate simulation time might be sufficient to observe cryptic pocket opening. Hence, we launched 320 100 ns-long independent simulations from an *apo* crystal structure of PM II (PDB 1LF4^25^).

To our surprise, we find that 32 microseconds of MD simulations do not reveal cryptic pocket opening in PM II. For the PM II cryptic pocket to open, three separate events must occur: Trp41 must change its sidechain orientation, Tyr77 must flip along χ-1, and the ‘flap’ domain must move away from the active site. Our *apo* seeded simulations sample both Tyr77 flipping and flap domain movement. However, we do not sample the change in Trp41 sidechain orientation (the distance between Trp41’s sidechain and the C-alpha of K72 remains large as seen in Fig. 3C). Hence, we conclude that PM II’s cryptic pocket opening is not captured with conventional MD simulations, though it is possible that large increases in the amount of sampling could enable us to observe Trp41 ring flipping that is necessary for cryptic pocket opening.

**Figure 3.**
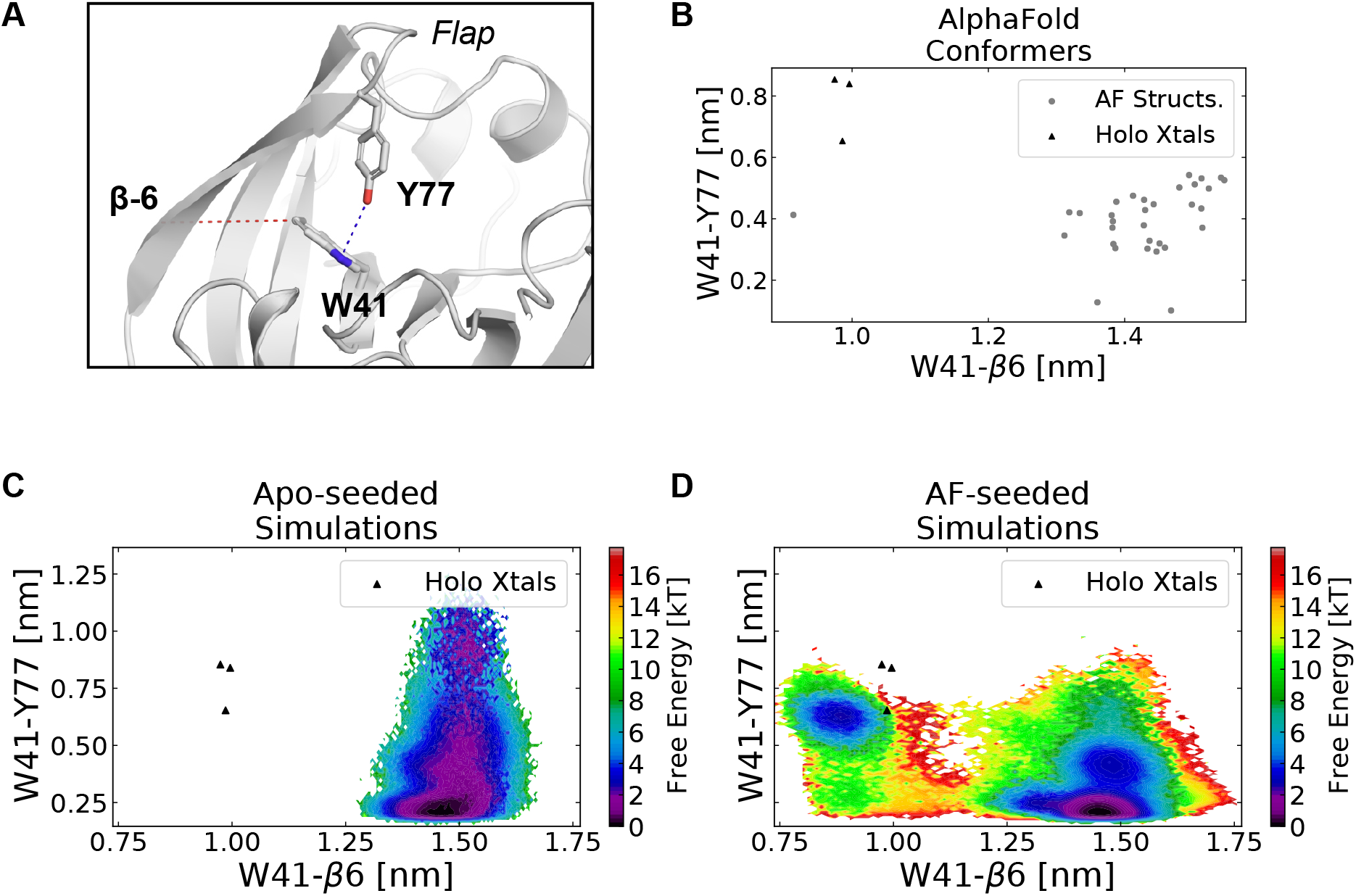
Launching simulations from AlphaFold-generated structures improves sampling of cryptic pocket opening in Plasmepsin II. **A)** Structure of PM II’s flap domain showing key residues involved in PM II’s cryptic pocket. Trp41 and Tyr77, part of the flap domain, are shown in sticks. We use the distances indicated in dotted lines to capture pocket opening. Specifically, the cryptic pocket is open when the minimum distance between Y77 and W41 is large (indicated with blue line) and the distance between the W41 sidechain (either atom CZ3 or CH2 depending on which is closer) and a reference residue in the 6^th^ beta-sheet (K72) is small (indicated with red line). **B)** Pocket distances for a set of 32 AlphaFold-generated conformers (grey dots) and *holo* crystal structures (black triangles) show that the AlphaFold ensemble includes partially open states for PM II. Trp41 is in its *holo* orientation in one of the AlphaFold structures, but the distance between Trp41 and Tyr77 is smaller than it is in *holo* crystal structures. **C)** A free energy surface from a Markov State Model from *apo*-seeded simulations shows that these simulations do not sample cryptic pocket opening. Though the flap dissociates as indicated by large Trp41-Tyr77 distances, Trp41 does not adopt the *holo* orientation, despite 32 microseconds of sampling. **D)** A free energy surface from a Markov State Model generated from AlphaFold-seeded simulations shows robust sampling of the open state. Both requirements for cryptic pocket opening are fulfilled as indicated by the overlay of *holo* crystal structures (black triangles) on the free energy surface.

### Seeding with AlphaFold accelerates exploration of the free energy landscape of PM II’s cryptic pocket

Given that the AF ensemble of PM II included diverse partially open structures (Fig. 3B)., we wondered if launching simulations from these structures would accelerate sampling of full cryptic pocket opening. The AF ensemble contains structures with different Trp41 orientations, including one with the Trp41 in the same orientation as *holo* crystal structures (Fig. S13A). Given that flap domain movement was sampled in the simulations initiated from the crystal structure, we hypothesized that we would observe open states in our simulations. We launched ten independent simulations of 100 ns in length for each of the 32 AlphaFold-generated starting structures (32 microseconds of aggregate simulation time). We also performed metadynamics simulations to generate a free energy landscape of Trp41 sidechain orientations using an orthogonal technique that could be compared against our unperturbed simulations.

We find that simulations launched from the AF ensemble sample cryptic pocket opening. Unlike in single-seeded simulations, we sample all three events required for cryptic pocket opening when simulations are launched from the AF ensemble (Fig. 3D). The Trp41 adopts a *holo*-like orientation while the distance between Trp41 and Tyr77 is large, creating a cavity for ligands to bind. Furthermore, we can build Markov State Models^26,40,41^ (MSMs) of the cryptic pocket ensemble to measure the probability of cryptic pocket opening. MSM are network models of free energy landscapes composed of many conformational states and the probabilities of transitioning between these states. Specifically, we constructed a MSM using a time-structure independent component analysis (tICA) projection of the backbone and χ-1 dihedrals within the cryptic pocket (see Methods). Despite starting from different starting structures, we find that our model is fully connected in this feature space, and we predict that the probability of cryptic pocket opening is 0.07, indicating that open states are a rare but non-negligible part of the ensemble.

Additionally, reasonable agreement between multiple simulation techniques suggests we have converged to the correct thermodynamics for the force field (Fig. 4). We use our MSM to construct a free energy landscape in the space of Trp41 χ-1 and χ-2 dihedral angles and compare against the free energy landscape generated by well-tempered metadynamics (see Methods). Overall, the two free energy landscapes identify similar free energy minima (Fig. 4). The deepest well in both landscapes corresponds to the Trp41 sidechain orientation seen in ligand-free structures (Fig. 4, S13B). There are minor differences in the two free energy landscapes with metadynamics predicting that the well centered on (−1, −2) is more probable than the MSM does. Furthermore, in metadynamics simulations the probability of the *holo* Trp41 orientation is 0.30 while in the MSM it is 0.08. Nonetheless, both methods predict that the flipped state necessary for pocket opening is a minor part of the ensemble.

**Figure 4.**
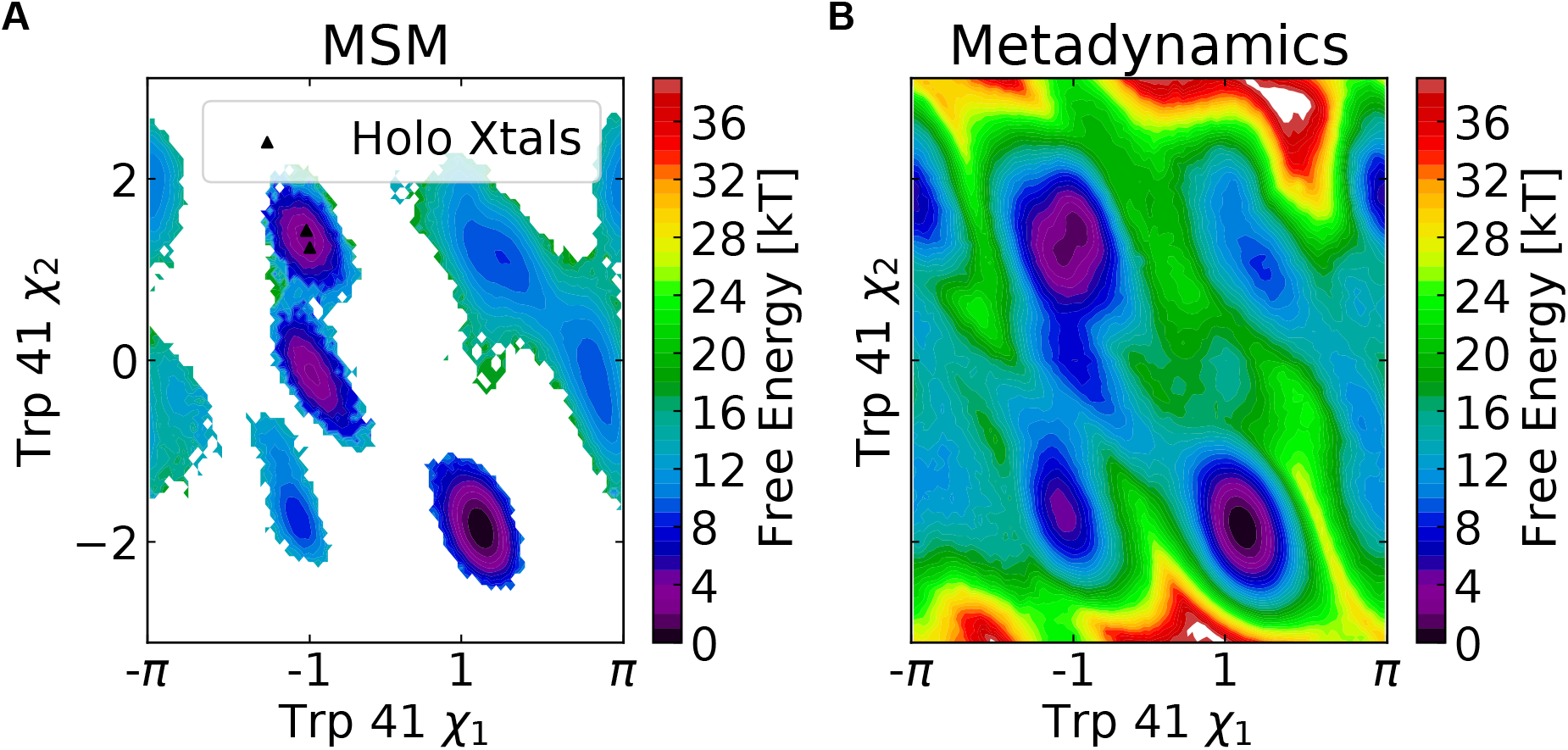
A Markov State Model built from AlphaFold-seeded simulations and metadynamics simulations yield similar free energy landscapes for plasmepsin II cryptic pocket opening. **A)** Free energy surface for Trp41 sidechain orientations derived from a Markov State Model constructed using dihedrals in the PM II cryptic pocket. *Holo* crystal structures are indicated with black triangles (three points are plotted though only two are visible). *Apo* crystal structures sample the well centered near (1, −2). **B)** Free energy surface from well-tempered metadynamics simulations using Trp41 χ-1and χ-2 angles as collective variables.

## Discussion

Certain cryptic pocket opening events remain difficult to sample with classical molecular dynamics simulations. As in previous work^12,13^, MD simulations launched from an *apo* PM-II structure failed to sample full cryptic pocket opening, even with an aggregate simulation time of 32 microseconds. This result makes PM-II an exception to a general trend. We have found that many cryptic pockets can be discovered with a handful of simulations of intermediate length (i.e., 40 ns)^1^. Furthermore, significant progress has been made in developing algorithms for cryptic pocket discovery, including Markov State Models^6,41^, enhanced sampling strategies like SWISH^42^, or adaptive sampling approaches like FAST^43^. However, we have previously seen that even adaptive sampling strategies can fail to sample known cryptic pockets. This likely stems from the difficulty of sampling rare events in classical molecular dynamics simulations.

The sampling strategy proposed here expands the available computational toolkit for cryptic pocket discovery and characterization without perturbing the underlying energy landscape (Fig. 1). Specifically, when assessing a protein as a drug target, we suggest generating diverse conformers of that protein by iteratively passing a stochastically subsampled multiple sequence alignment to AlphaFold. Next, we propose using pocket detection tools, such as LIGSITE^44^, fpocket^45^, or P2rank^46^, to identify pockets that are absent in *apo* experimental structures or an AlphaFold-predicted structure using a complete MSA. In some cases, this will be sufficient to uncover novel cryptic pockets (Fig. 2A). However, if this approach yields partial opening or one is interested in assessing the equilibrium probability of a cryptic pocket opening, we propose using molecular dynamics simulations followed by Markov State Model construction. As demonstrated here with PM II, this strategy can greatly accelerate the discovery and characterization of cryptic pockets.

Drug discovery efforts directed towards plasmepsins illustrate that targeting cryptic pockets is a generally promising strategy for discovering selective and potent inhibitors. Ligands that bind at the PM II cryptic site have enhanced potency and selectivity towards PM II compared with other plasmepsins from *Plasmodium falciparum* (Fig. S16A and S16B). Furthermore, ligands that bind in the cryptic pockets do not inhibit human pepsin-like aspartic proteases (e.g., pepsin, cathepsin D and E)^29^. To further illustrate the utility of targeting the PM-II cryptic site, we used metadynamics to compare the unbinding of an inhibitor from the cryptic site with the unbinding of a ligand from the active site (Fig. S16). We find that the ligand which binds at the cryptic pocket has a ~25 kJ/mol higher free energy barrier to unbinding because the tyrosine in the cryptic pocket acts as a lid over the ligand (Fig. S16C). Slower unbinding kinetics may explain why ligands that bind in the PM II cryptic pocket are more potent and selective. We expect these same principles will apply in other systems.

## Conclusions

We have demonstrated that AlphaFold can be used to accelerate the discovery and characterization of cryptic pockets. When its input multiple sequence alignment is stochastically subsampled, AlphaFold generates diverse conformers of proteins known to form cryptic pockets. In 6 out of 10 examples of proteins known to form cryptic pockets, AlphaFold samples the open state (Fig. 2A). Impressively, AlphaFold also makes predictions of the open state even when the *holo* structure was deposited after AlphaFold was trained. In other cases, like with plasmepsin II, AlphaFold samples partially open states (Fig. 2C). For example, in plasmepsin II, the ensemble of AF structures includes structures with a tryptophan sidechain in its *holo* orientation, even though 32 microseconds of MD simulations launched form an apo crystal structure do not sample tryptophan flipping. We find that launching simulations from this ensemble accelerates sampling of the open state (Fig. 3). Furthermore, because we observe both pocket opening and closing events, we can use a Markov State Model to generate a free energy landscape of pocket conformations that is in reasonable agreement with a similar landscape generated from metadynamics. Thus, we propose an efficient strategy to discover cryptic pockets that we hope becomes indispensable to future structure-based drug design efforts.

## Acknowledgements

This work was funded by NSF MCB 2218156, NIH NIA RF1AG067194, and NIH NIGMS R01GM124007. G.R.B. holds a Packard Fellowship from the David and Lucile Packard Foundation. AM was supported by the National Institutes of Health F30 Fellowship (1F30HL162431-01A1).

## Supporting Information

All input files and analysis scripts corresponding to this study can be accessed here: https://github.com/sbhakat/AF-cryptic-pocket.

