## Supplement for "Accelerating cryptic pocket discovery using AlphaFold"

**Supporting Information Available**

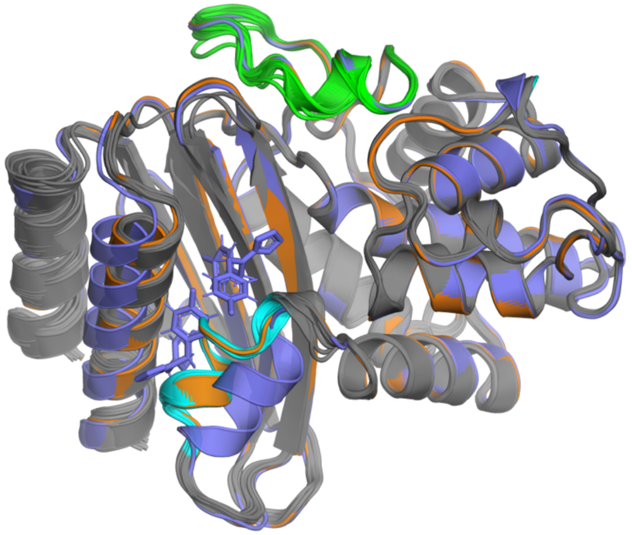

**Figure S1:** The ensemble of TEM beta lactamase structures from AlphaFold does not include conformations where the Horn or omega pockets are open. The Horn (cyan) and omega loop (green) from the 32-model AlphaFold ensemble (gray) do not sample the open conformation seen in the *holo* structure (PDB: 1PZO, purple) and show limited deviation from the *apo* structure (PDB: 1JWP, orange).

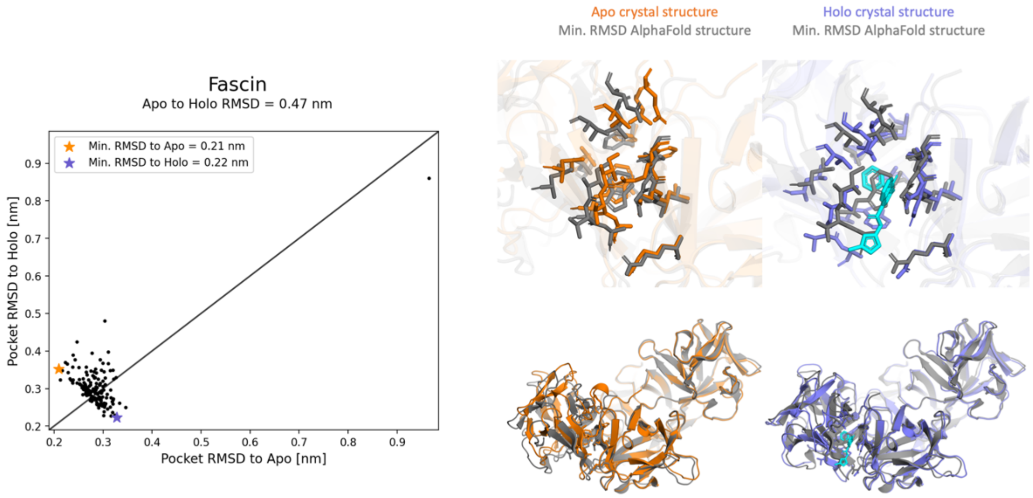


**Figure S2:** The RMSD of pocket residues of the AlphaFold structures to the *apo* structure (PDB 3P53, orange) and the *holo* structure (PDB 6i11, purple) for fascin. The RMSD between *apo* and *holo* structures is 0.47 nm. The structures on the right display the AlphaFold structures (gray) closest to either the *apo* or *holo* crystal structure, with their RMSD values depicted as stars on the scatterplot.

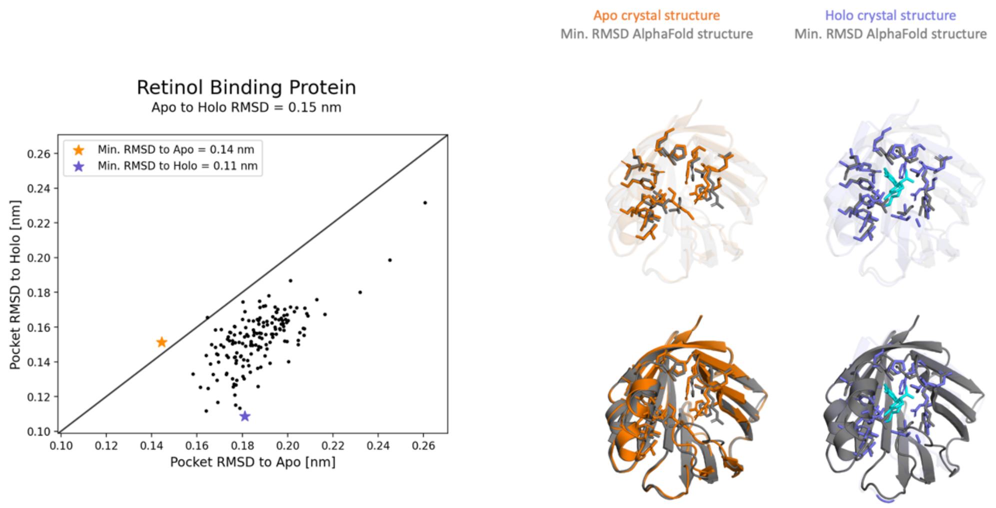


**Figure S3:** The RMSD of pocket residues of the AlphaFold structures to the *apo* structure (PDB 5H9A, orange) and the *holo* structure (PDB 6E51, purple) for retinol binding protein 1. The RMSD between *apo* and *holo* structures is 0.15 nm. The structures on the right display the AlphaFold structures (gray) closest to either the *apo* or *holo* crystal structure, with their RMSD values depicted as stars on the scatterplot.

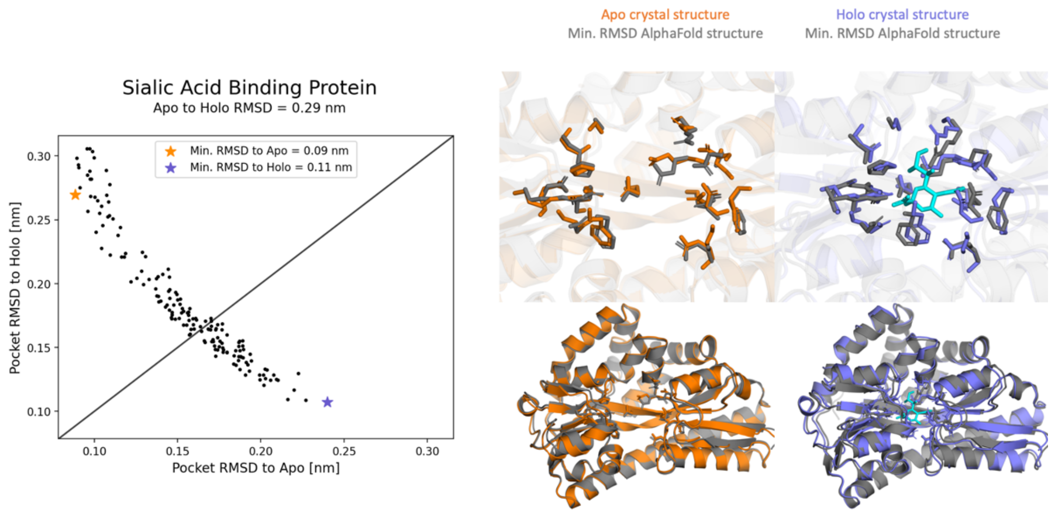


**Figure S4:** The RMSD of pocket residues of the AlphaFold structures to the *apo* structure (PDB 2CEY, orange) and the *holo* structure (PDB 6H76, purple) for the sialic acid binding protein from *H. influenzae*. The RMSD between *apo* and *holo* structures is 0.29 nm. The structures on the right display the AlphaFold structures (gray) closest to either the *apo* or *holo* crystal structure, with their RMSD values depicted as stars on the scatterplot.

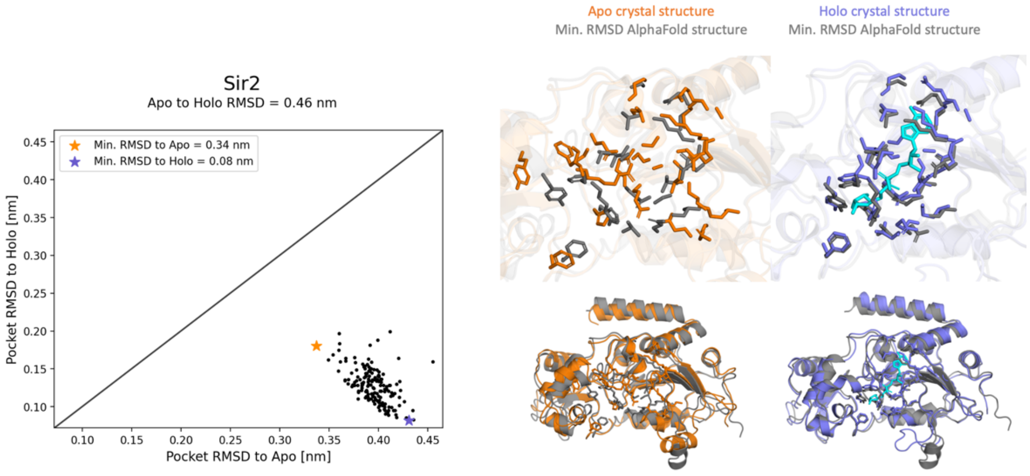


**Figure S5:** The RMSD of pocket residues of the AlphaFold structures to the *apo* structure (PDB, orange) and the *holo* structure (PDB, purple) for Sir2. The RMSD between *apo* and *holo* structures is 0.46 nm. The structures on the right display the AlphaFold structures (gray) closest to either the *apo* or *holo* crystal structure, with their RMSD values depicted as stars on the scatterplot.

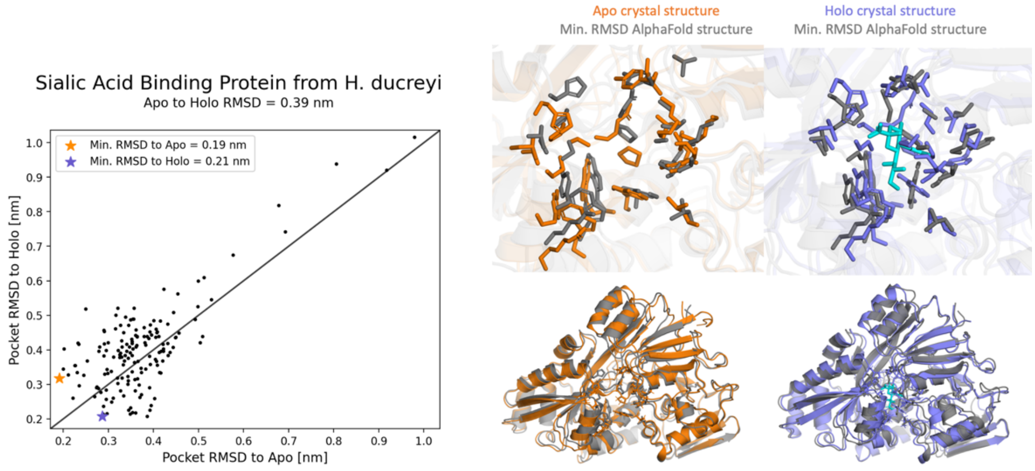


**Figure S6:** The RMSD of pocket residues of the AlphaFold structures to the *apo* structure (PDB 5Z4A, orange) and the *holo* structure (PDB 5YYB, purple) for sialic acid binding protein from *H. duceryi*. The RMSD between *apo* and *holo* structures is 0.39 nm. The structures on the right display the AlphaFold structures (gray) closest to either the *apo* or *holo* crystal structure, with their RMSD values depicted as stars on the scatterplot.

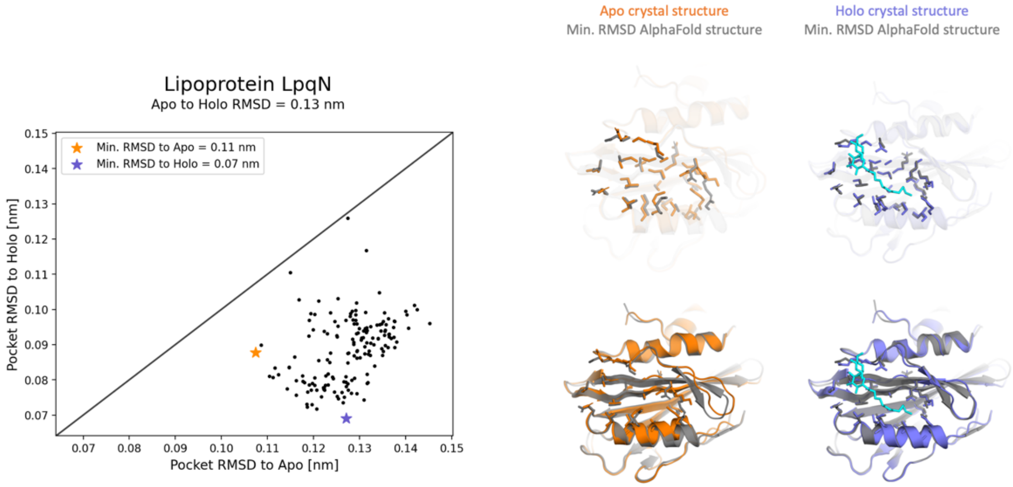


**Figure S7:** The RMSD of pocket residues of the AlphaFold structures to the *apo* structure (PDB 6E5D, orange) and the *holo* structure (PDB 6E5F, purple) for lipoprotein LpqN. The RMSD between *apo* and *holo* structures is 0.13 nm. The structures on the right display the AlphaFold structures (gray) closest to either the *apo* or *holo* crystal structure, with their RMSD values depicted as stars on the scatterplot.

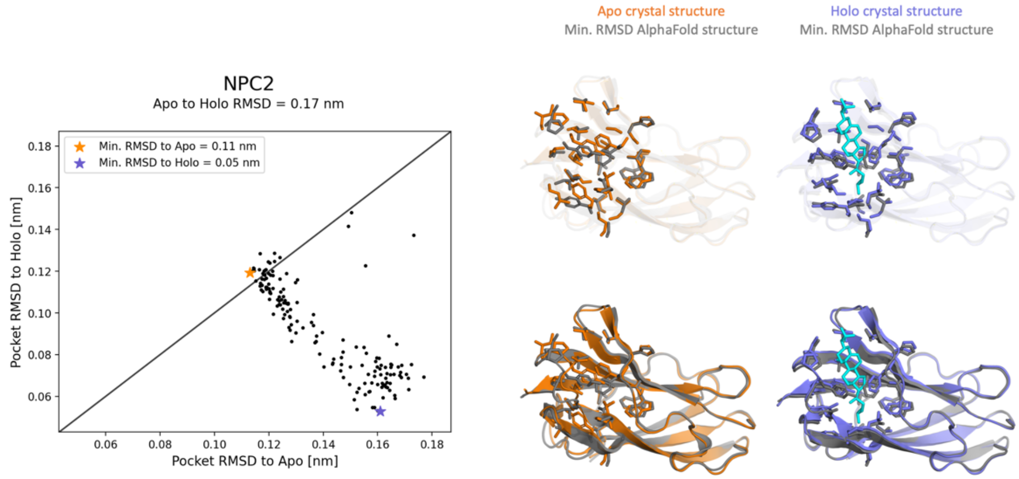


**Figure S8:** The RMSD of pocket residues of the AlphaFold structures to the *apo* structure (PDB 1NEP, orange) and the *holo* structure (PDB 2HKA, purple) for NPC2. The RMSD between *apo* and *holo* structures is 0.17 nm. The structures on the right display the AlphaFold structures (gray) closest to either the *apo* or *holo* crystal structure, with their RMSD values depicted as stars on the scatterplot.

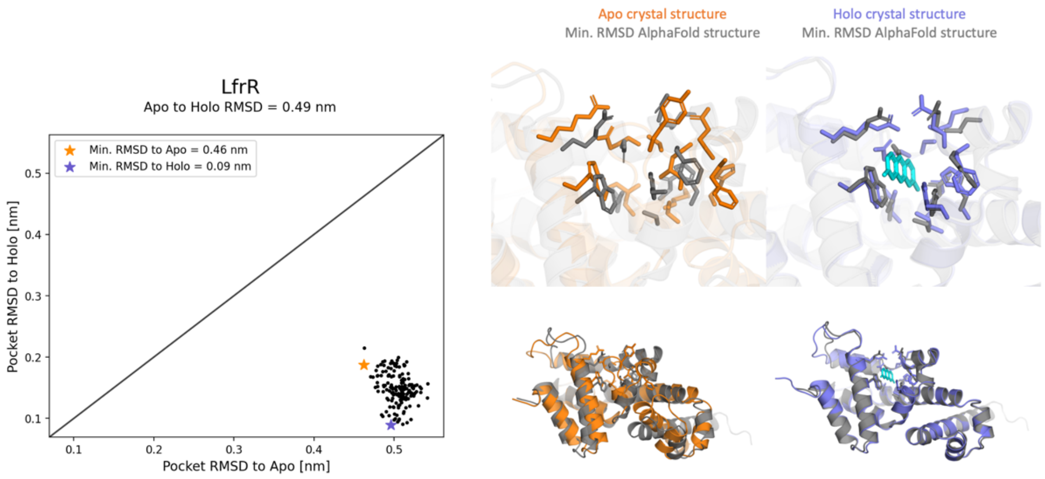


**Figure S9:** The RMSD of pocket residues of the AlphaFold structures to the *apo* structure (PDB, orange) and the *holo* structure (PDB, purple) for LfrR. The RMSD between *apo* and *holo* structures is 0.49 nm. The structures on the right display the AlphaFold structures (gray) closest to either the *apo* or *holo* crystal structure, with their RMSD values depicted as stars on the scatterplot.

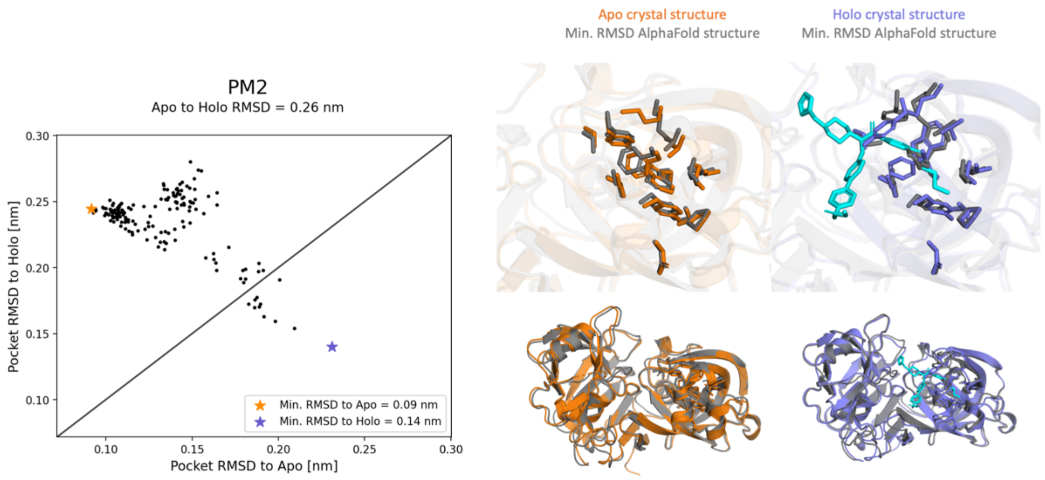


**Figure S10:** The RMSD of pocket residues of the AlphaFold structures to the *apo* structure (PDB 1LF4, orange) and the *holo* structure (PDB 2BJU, purple) for Plasmepsin II. The RMSD between *apo* and *holo* structures is 0.26 nm. The structures on the right display the AlphaFold structures (gray) closest to either the *apo* or *holo* crystal structure, with their RMSD values depicted as stars on the scatterplot.

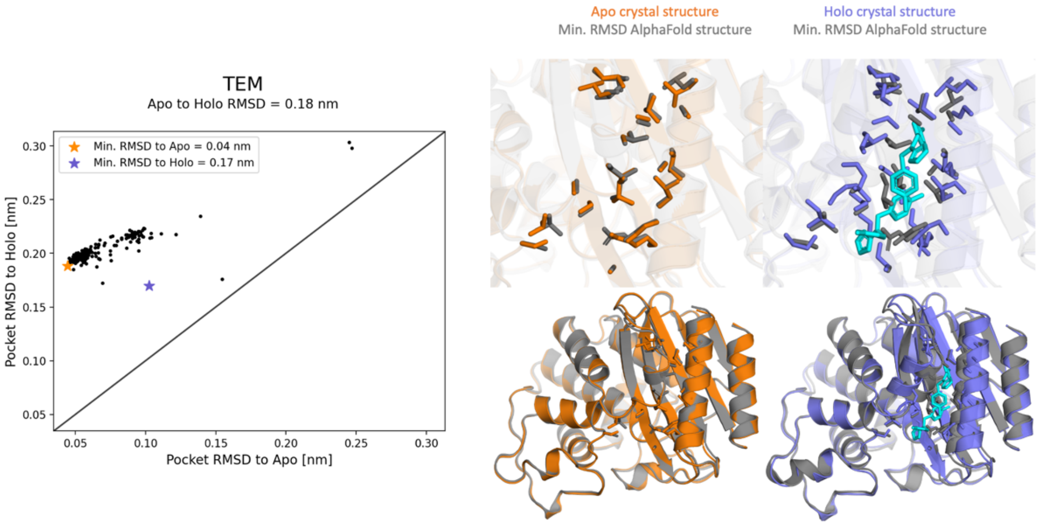


**Figure S11:** The RMSD of pocket residues of the AlphaFold structures to the *apo* structure (PDB 1JWP, orange) and the *holo* structure (PDB 1PZO, purple) for TEM beta-lactamase. The RMSD between *apo* and *holo* structures is 0.18 nm. The structures on the right display the AlphaFold structures (gray) closest to either the *apo* or *holo* crystal structure, with their RMSD values depicted as stars on the scatterplot.

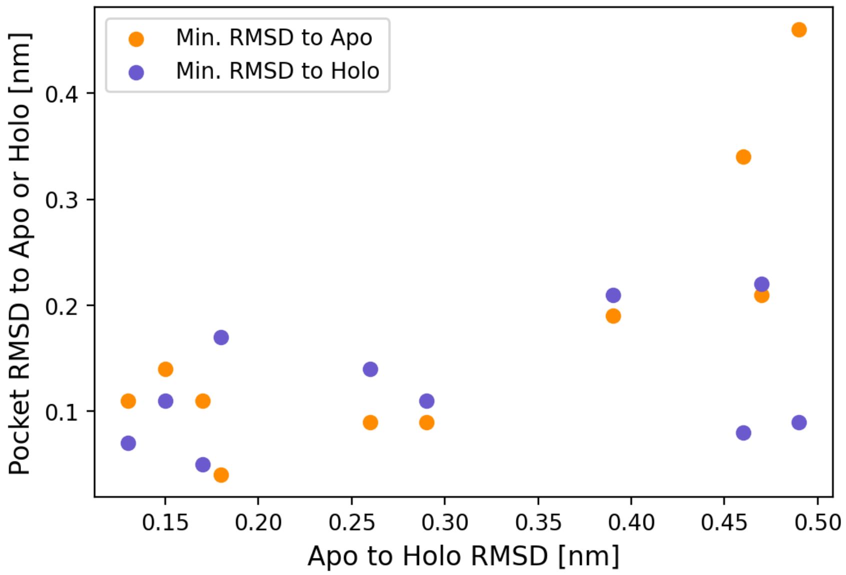


**Figure S12:** Correlation between the RMSD of *apo* to *holo* structures and the minimum RMSD of the AlphaFold structures to the *apo* (purple) or *holo* (orange) structures.

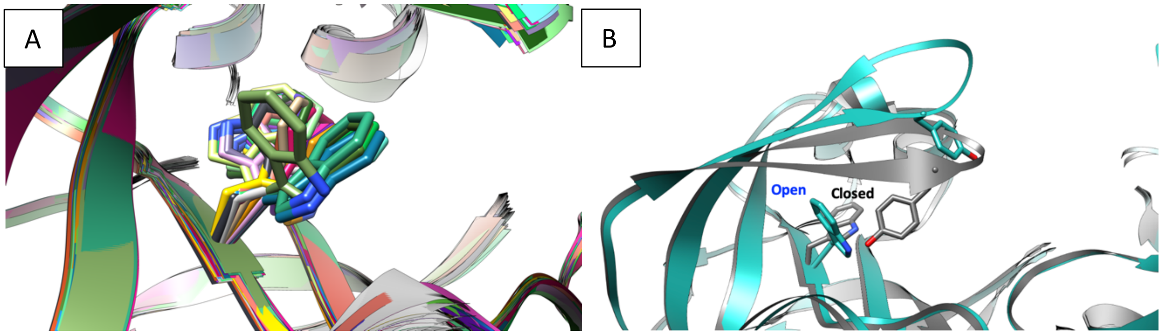


**Figure S13:** Stochastic subsampling of its input MSA allows AlphaFold to generate a diverse ensemble of Trp41 conformations, as shown in A. The differences in Trp41 orientation between *apo* (PDB: 1LF4, grey) and *holo* PM II structures (PDB: 2BJU, blue) with an open cryptic pocket are shown in B.

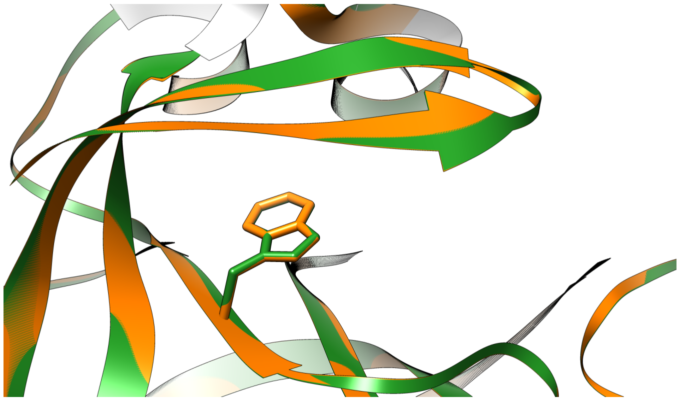

**Figure S14:** The *apo* PM II structures (orange: 1LF4, green: 3F9Q) show the Trp41 in a conformation that blocks access to the cryptic site.

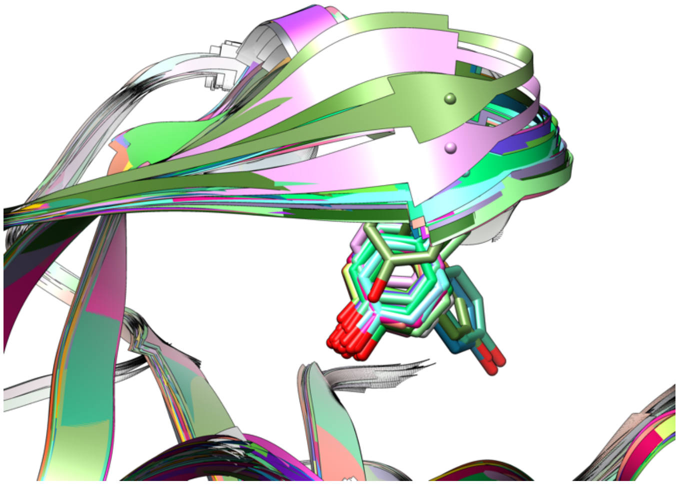

**Figure S15:** The AlphaFold generated ensemble of PM II structures samples the Tyr77 conformation from the *holo* structure.

A

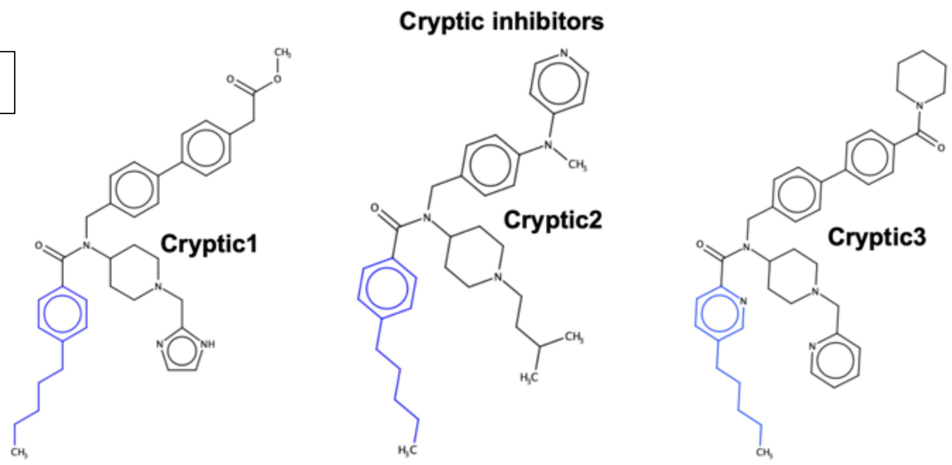


B

| | PM I | PM II | PM IV |
|---|---|---|---|
| **Cryptic1** | N/A | **0.034** | N/A |
| Cryptic2 | 1.123 | **0.113** | 19.440 |
| Cryptic3 | 0.711 | **0.054** | 3.214 |
| **Traditional** | 4.1 | **0.8** | 0.25 |

C

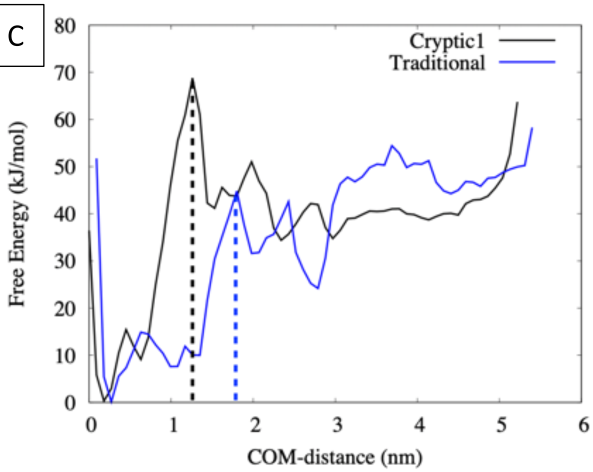


D

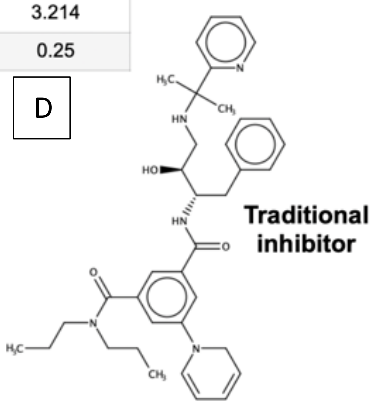


**Figure S16:** PM II inhibitors which bind the cryptic pocket have enhanced selectivity and a larger barrier to unbinding. (A) IC50 values (in micro-molar) highlight selectivity of cryptic pocket-binding inhibitors compared to an active site inhibitor. (B) PM II inhibitors which bind in the cryptic pocket have a higher free energy barrier to unbinding. Reweighted free energy surface along unbinding distance (COM-distance) for PM II-inhibitor complexes shows a ~ 25kJ/mol higher free energy barrier associated with an inhibitor that binds in the cryptic pocket (PDB: 2BJU) as compared to active site inhibitor (PDB: 4YA8). Black and blue lines showing the apparent free energy barrier in each case.

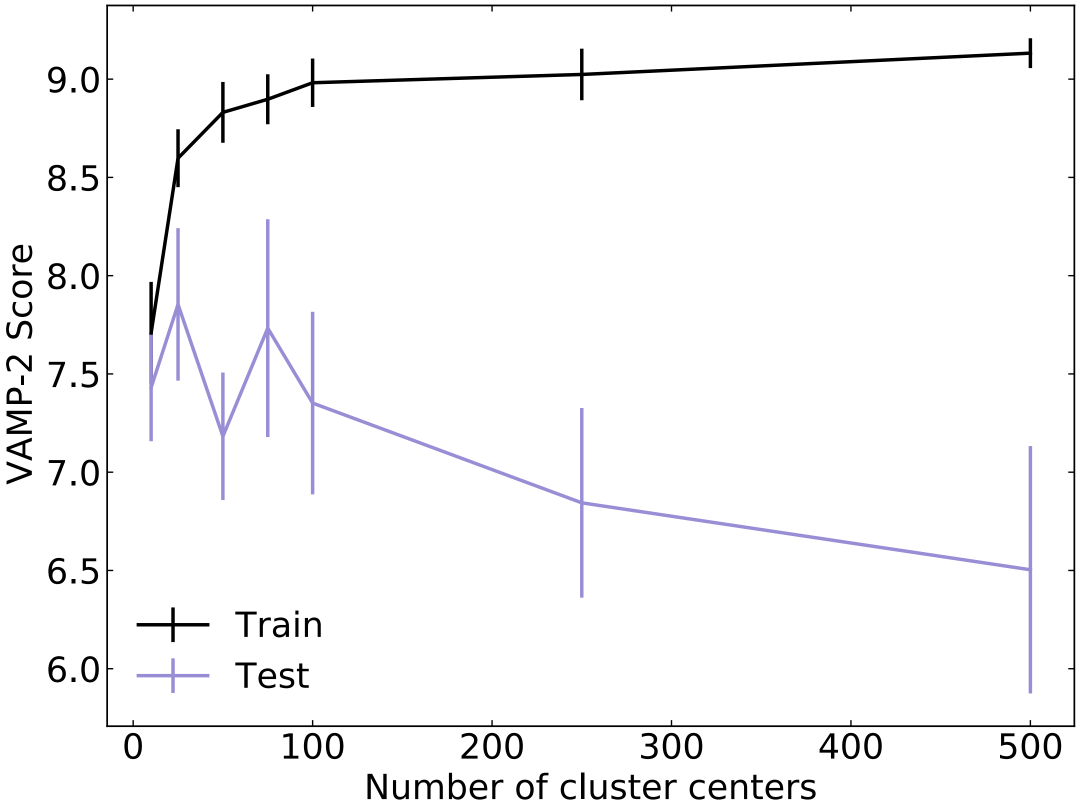


**Figure S17:** VAMP-2 scores across 10 trials for the AF-seeded ensemble shows the highest mean test set score is achieved when MSMs contain 25 microstates.

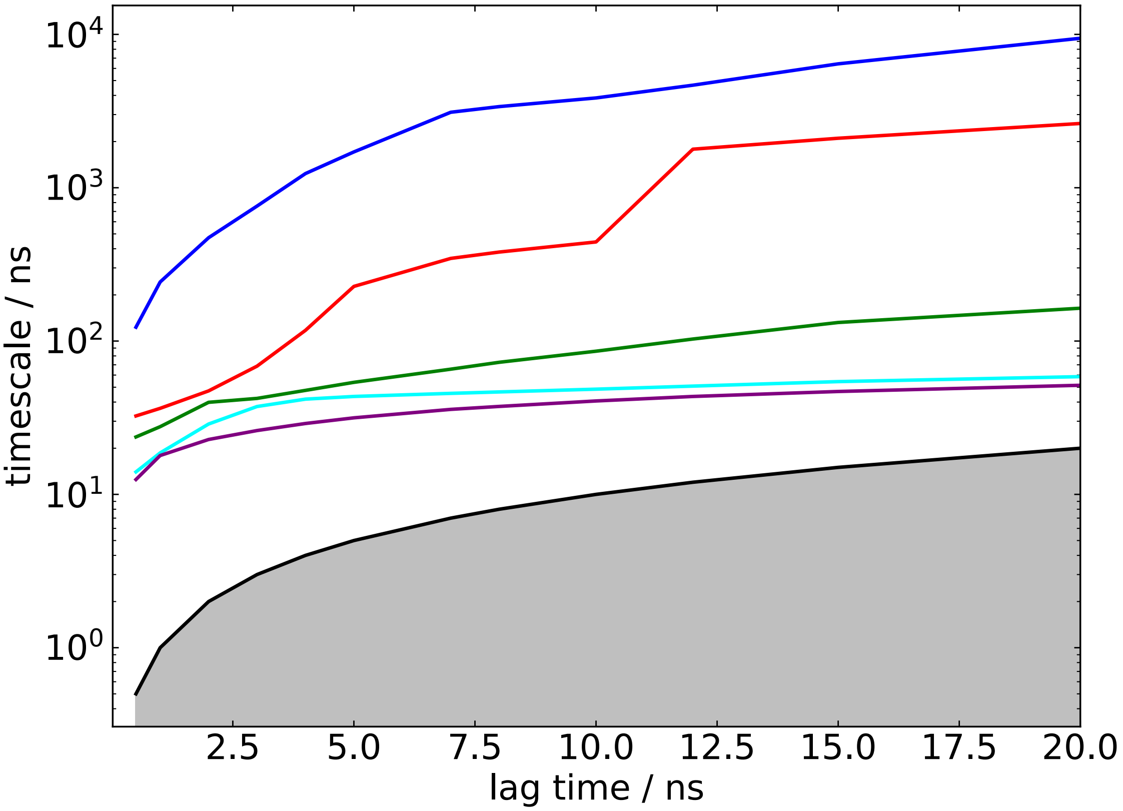


**Figure S18:** Implied timescales plot for the *apo*-seeded simulation dataset shows that first 5 timescales converge around 12 ns on a logarithmic scale.

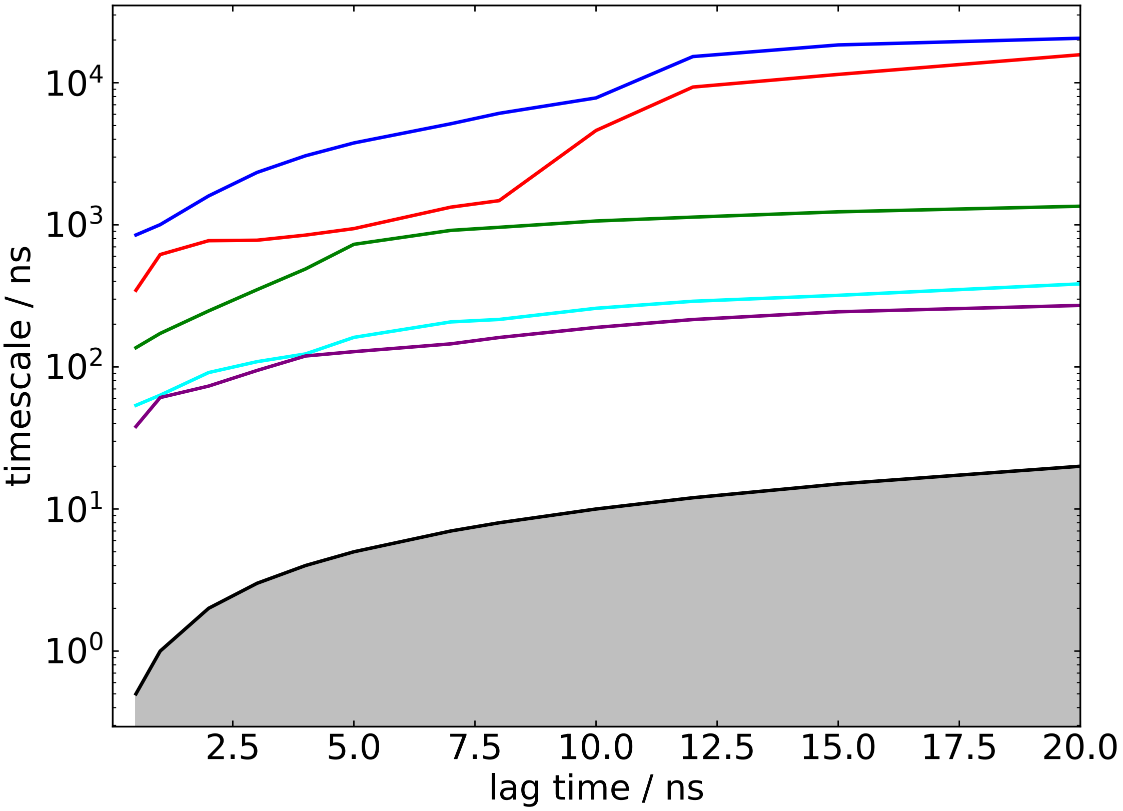


**Figure S19:** Implied timescales plot for the AF-seeded simulation dataset shows that first 5 timescales similarly converge around 12 ns on a logarithmic scale.
